# Multiplexed selectivity screening of anti-GPCR antibodies

**DOI:** 10.1101/2022.11.24.517810

**Authors:** Leo Dahl, Ilana B. Kotliar, Annika Bendes, Tea Dodig-Crnković, Samuel Fromm, Arne Elofsson, Mathias Uhlén, Thomas P. Sakmar, Jochen M. Schwenk

## Abstract

G protein-coupled receptors (GPCRs) control critical cellular signaling pathways. Therapeutic agents, such as antibodies (Abs), are being developed to modulate GPCR signaling pathways. However, validating the selectivity of anti-GPCR Abs is challenging due to sequence similarities of individual receptors within GPCR subfamilies. To address this, we developed a multiplexed immunoassay to test >400 anti-GPCR Abs from the Human Protein Atlas targeting a customized library of 215 expressed and solubilized GPCRs representing all GPCR subfamilies. We found that ~61% of Abs were selective for their intended target, ~11% to bind off-target, and ~28% not to bind any GPCR. Antigens of on-target Abs were, on average, significantly longer, more disordered, and less likely to be buried in the interior of the GPCR protein than the other Abs. These results provide important insights into the immunogenicity of GPCR epitopes and form a basis for the design of therapeutic Abs and the detection of pathological auto-antibodies.

**TEASER:** A multiplexed library-to-library selectivity analysis of 400 anti-GPCR antibodies within subfamilies of 200 solubilized receptors.

## INTRODUCTION

The superfamily of G protein-coupled receptors (GPCRs) comprises ~ 750 membrane proteins that mediate signaling pathways for many aspects of cellular physiology and intercellular communication. GPCRs are also targets for ~ 30% of approved therapeutic drugs (*1, 2*). Disruptions in GPCR function or signaling contribute to the pathophysiology of numerous disease states. Even though GPCRs are one of the most targeted protein families in pharmaceutical drug development, robust and high-throughput methods to study these dynamic membrane proteins remain limited (*3*).

Among the possible approaches to studying membrane proteins, antibodies (Abs) have been used in various bioanalytical applications. In recent years, however, the perception of the omni-applicability of Abs has drastically changed (*4*). In fact, the research community has called for an end to using unvalidated Abs as an off-the-shelf solution to study human biology (*5*). This call led to the creation of guidelines for Ab validation (*6*) and database registration to track their use across different methods and sample types (*7*). A key outcome of these efforts was recognizing that achieving better data from immuno-capture assays requires defining Ab performance and utility according to sample type, target preparation, and analytical platform (*8*). Therefore, dedicated and systematic Ab validation efforts have been conducted for several technology platforms and assay formats, including immunohistochemistry of tissue samples (*9*), cellular immunofluorescence assays (*10, 11*), and Western immunoblots of sample lysates (*12, 13*) and of soluble proteins found in blood (*14*). Defining selective Abs validated for specific applications also facilitates the identification of specific Ab epitopes to measure possible effects from drugs developed against a target of interest.

To evaluate the selectivity and potential utility of existing Abs generated against GPCR targets, we established a multiplexed approach in which a large library of anti-GPCR Abs was tested in parallel against a large library of engineered GPCRs. We build upon a previous proof-of-concept study that employed a suspension bead array (SBA) assay and a set of heterologously expressed and solubilized GPCR constructs that included genetically-encoded monoclonal Ab (mAb) epitope tags (*15*). Each multiplexed SBA assay exposes all Abs to in the library set to one GPCR at a time. Each GPCR is thereby serves as the on-target reactivity of a subset of the Abs, while simultaneously serving as a probable off-target for all the other Abs in the SBA. With the objective to challenge the selectivity of each Ab, we increase the likelihood of off-target GPCR recognition by presenting overexpressed off-targets and in the absence of the intended GPCR. We found that 248 of 407 anti-GPCR Abs tested against a library of 215 GPCRs were selective for their intended targets. To provide a transparent view of the data set, we also present a webbased platform that enables data browsing and analysis. Our results provide insights concerning the immunogenicity of GPCRs, and for the design and application of anti-GPCR Abs as tool reagents and therapeutic agents.

## RESULTS

We established a multiplexed framework to interrogate the selectivity of Abs raised against GPCRs. The procedure combined parallel expression of GPCRs with multiplexed protein capture assays. The detection of the immunocaptured proteins is assured via engineered epitope tags that allow the determination of relative quantities of the GPCRs in solution. The data are integrated into a web-based interface to browse the selectivity of each Ab and GPCR (**Fig. 1**).

**Fig. 1.**
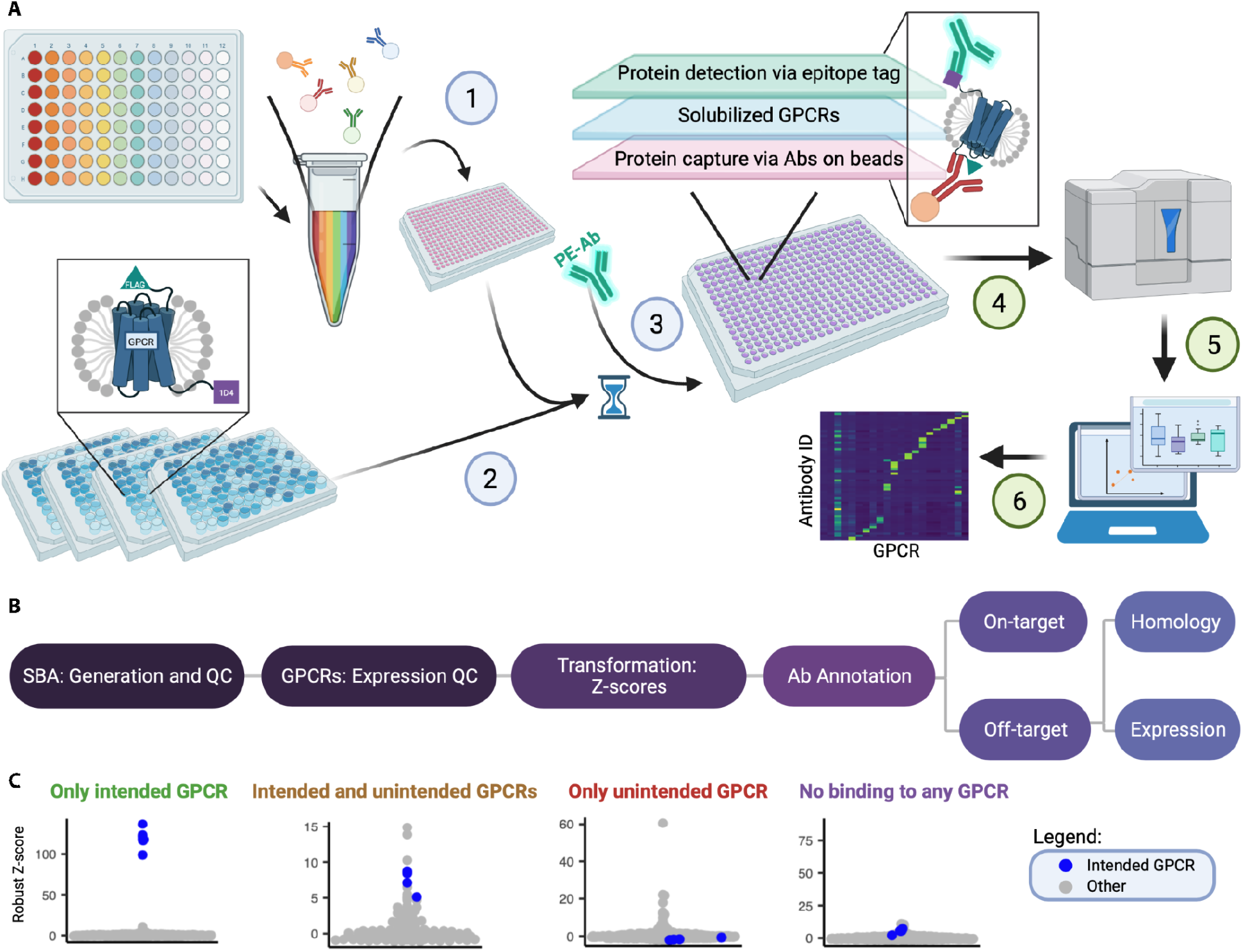
Experimental workflow and data analysis. (**A**) An SBA assay experimental workflow was adapted from previous work (*15*). The numbers within the schematic indicate the following steps: 1 - unique Abs were coupled to beads to create SBAs. Beads were then grouped by the phylogenetic subfamily of their GPCR target and pooled to generate six SBAs with 12 to 126 different populations of capture beads per pool. Bead pools were dispensed into 384-well plates for the assay. 2 - dual epitope-tagged GPCRs were expressed in Expi293F cells. The cell membranes were then solubilized in detergent, which resulted in heterogeneous mixtures of solubilized membrane proteins, including the GPCRs. Total protein concentration was normalized across samples, and aliquots were transferred to the assay plate containing the SBAs. Hourglass - the assay plate was incubated overnight. 3 - PE-conjugated detection mAb targeting the 1D4 tag was then added. 4 - a Luminex FlexMap 3D instrument was used to measure the reporter fluorescence while simultaneously reading the barcode of each bead. 5 - the data was used to create an interactive web interface. 6 - the specificity and selectivity of the tested Abs was determined. (**B**) Flowchart of data analysis. Data generated as described in (**A**) was subject to quality control (QC) tests for SBA generation and GPCR expression. Next, the data were scaled and centered. Abs tested were then annotated as binding “On-target” or “Off-target”, and the latter were further sub-classified by the proposed cause of Off-target binding (homology or expression). (**C**) Schematic of data visualization. Each example beeswarm plot represents a single Ab, and each dot represents a single cell-based sample. The y-axis is the Robust Z-score, and there is no quantitation on the X-axis. The plots enable quantitative identification of four types of Ab binding behavior. The blue dots represent samples that ectopically express the target GPCR, and the gray dots represent samples that ectopically express GPCRs other than the target. Abs binding only the intended GPCR (green label) are considered validated. Abs that bind intended and unintended GPCR targets (ochre label) or unintended GPCRs only (red label) are analyzed further to identify a potential root cause. Abs that do not bind to any target (purple label) are considered not validated. Created in BioRender.com

We tested 407 polyclonal Abs (pAbs), of which 399 Abs were developed in the Human Protein Atlas (HPA) project and eight Abs were from other commercial sources (CABs). The HPA Abs were raised against recombinant protein fragments of 50-150 amino acids (aa) in length, which were selected using a bioinformatic algorithm (*16*). The approach aims to select unique primary structures, and to avoid transmembrane regions and amino acid sequences with homology to any of the other 20,000 protein-encoding gene products. Each HPA Ab was affinity purified on its respective antigen and then tested in a pipeline (*17*). Assays of HPA included initial Ab binding assays on protein arrays that were built on 384 recombinant antigens (*18*) as well as Western Blots containing tissue, cell, and plasma lysates (*19*) with six different types of samples. The primary utility of HPA Abs is to map the expression of the human protein across tissues and cells(*20, 21*). The tissues are interrogated as sections of paraffin-embedded tissues, while the fixated cells are analyzed for the subcellular location of the protein targets.

To generate our GPCR library, we chose Expi293F cells, which are suspension-adapted Human Embryonic Kidney (HEK) 293T cells that ectopically expressed each of the 215 dual epitope-tagged GPCRs (**Fig. S1**). Each GPCR construct includes an N-terminal FLAG tag and a C-terminal 1D4 tag, except for four GPCRs belonging to the Frizzed (FZD) subfamily, which have an N-terminal HA epitope tag and a C-terminal 1D4 epitope tag. The tags are used to compare the quantities of receptors added to the assay and as a detection system that is common for all GPCRs. To harvest the receptors, membranes from Expi293T cells were solubilized with dodecyl maltoside (DM, full name n-dodecyl-b-D-maltoside) detergent buffer. The detergent micelles that were generated contain a heterogenous mixture, including the overexpressed GPCRs from cell membranes. These samples were flash-frozen, stored, and thawed shortly before the analyses.

To create suspension bead arrays (SBAs), each Ab was covalently coupled to a distinct population of color-coded magnetic beads. Abs for a specific GPCR subfamily were then pooled into subfamily-specific SBAs (**Fig. 1A, Table S1**). The six subfamily groupings of the SBA correspond to the GRAFs classification system: rhodopsin (divided into alpha, beta, gamma and delta), “other” and glutamate, adhesion, secretin and FZD (GSAF) combined into one group. Each SBA also contains Abs against the engineered epitope tags to determine the expression of the GPCRs in each assay. Since each GPCR construct contains the same pair of epitope tags, the relative abundance of each GPCR was compared across replicated preparations, GPCR classes or subfamilies via capture of the FLAG tag, detection of the 1D4 tag, followed by a readout in a flow cytometer (**Fig. 1A, Fig. 2A-B**). We confirmed the agreement in expression between biological and technical replicates for three selected GPCRs from different phylogenetic groups **(Fig. 2C**). For the selectivity screen, we used one technical replicate of four biological replicates for each GPCR-containing sample. Technical duplicates were included for a subset of GPCRs chosen at random. An essential aspect of the experimental design was to test many Abs in parallel for all six GPCR subfamilies. With this design, each Ab is exposed to its intended and overexpressed on-target GPCR and all other overexpressed family members, representing the most probable off-target GPCRs. While the initial step is to assure the detection of the intended target of each Ab, the utility of each Ab increases if it does not bind to any other overexpressed off-target. Such an interrogation reveals which Abs are selective to recognize the solubilized receptors in multiplexed protein capture assays.

**Fig. 2.**
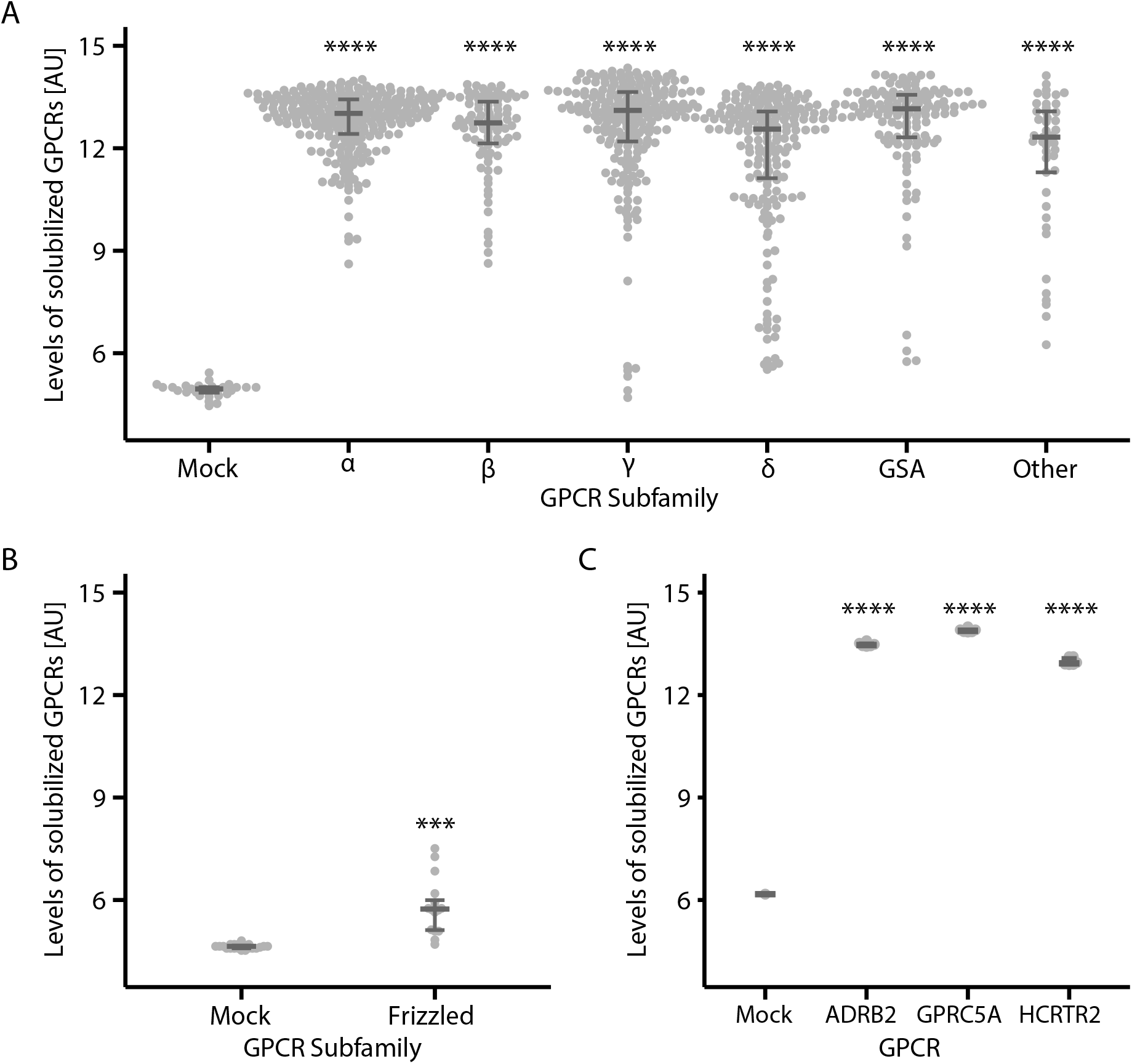
Quantification of overexpressed and solubilized GPCRs by phylogenetic group. **(A)** Relative quantities of solubilized dual epitope-tagged GPCRs were determined by SBA immunoassay. Here FLAG was used for capture and 1D4 for detection. GPCRs are grouped by subfamily, and expression data are plotted as log2 of median fluorescence intensity (MFI). **(B)** Relative quantities of solubilized dual epitope-tagged GPCRs in the frizzled subfamily. Here HA was used for capture and 1D4 for detection. Expression data are plotted as the log2 MFI. Horizontal bars represent the medians of each group with the 25^th^ and 75^th^ percentiles. **(C)** Representative examples of GPCR expression and solubilization reproducibility. GPCRs selected from three different subfamilies were expressed on three separate occasions (biological replicates), each quantified in technical duplicates (N=6). Quantification was carried out as described in **(A)**. Significance was determined by a one-way ANOVA (with p < 0.05) followed by Dunnett’s multiple comparison test to mock. Here, **** indicates a p < 0.0001, and *** stands for p < 0.001. Sample sizes and p-values are listed in **Table S2**.

To enable a systematic assessment of Ab selectivity data at scale, the median fluorescence intensity (MFI) levels recorded per bead population and sample were subjected to several quality control (QC) steps to ensure successful Ab-bead coupling and to quantify the amounts of solubilized GPCRs added to the assay (**Fig. 1B**). The raw data were then scaled, centered and transformed into Z-scores and robust Z-scores (R.Z-scores) for each anti-GPCR Ab separately. Finally, each Ab was annotated according to the exhibited on- and off-target selectivity. For off-target events, further analyses were conducted that considered sequence homology and expression levels as possible causes of the cross-reactivity. The performance of each Ab was visualized in beeswarm plots with R.Z-scores (for examples, see **Fig. 1C**). These examples illustrate each of the four types of Ab annotations: on-target, co-target, off-target or no-target. The beeswarm plots and other criteria for the investigated GPCRs and tested Abs are provided in detail in a browsable interface (*Shiny* App).

### Discovery of selective GPCR antibodies

Our study investigated the selective binding of 407 Abs against 205 GPCRs. Although there were 215 GPCRs in the library, Abs chosen against ten GPCRs did not pass quality control for the assay. These ten GPCRs were used as off-target candidates only.

At first, we checked the expression and solubilization efficiency of each GPCR subfamily member compared with the negative control (mock). As shown in **Fig. 2A-B** and **Table 1A**, most GPCRs, particularly those from the rhodopsin alpha and beta subfamilies, were present in sufficient amounts. Only a handful of solubilized GPCRs from the rhodopsin gamma and delta, GSAF and “other” subfamilies were detected at lower levels. The GPCRs assigned to the smaller FZD subfamily within the GSAF group expressed the lowest efficiently. Considering the levels of GPCRs allowed us to judge the capability of Abs for selective or cross-reactive binding to GPCRs.

**Table 1.**
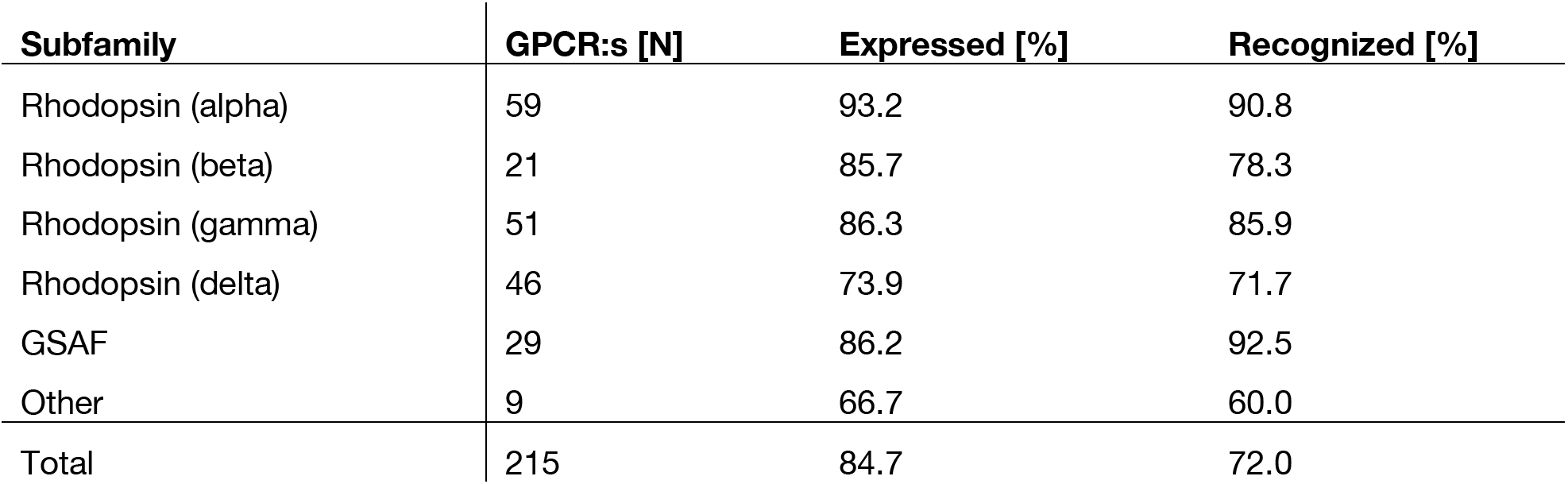
GPCR expression and antibody performance. Per subfamily summary of GPCRs (number of members, number expressed) and antibodies (validation) are given. **Table 1A**. The total number of generated GPCRs, the percentage of GPCRs with evidence of expression (determined via the epitope tags), and the percentage of expressed GPCRs recognized by the tested anti-GPCR Abs. The latter category is limited to only GPCRs targeted by at least one Ab.

To determine an appropriate cut-off for defining Ab selectivity, we constructed a model to test how many Abs would pass and recognize their intended GPCR or fail due to insufficient target enrichment. We used R.Z-scores and the density peaks of the population data obtained per Ab and applied a function of standard deviations (SDs) to determine the global selectivity threshold (**Fig. 3A**). We selected a cut-off of 12x SD over the density peak, as this SD level coincided with the peak in on-target recognitions and the beginning of the plateaus for the on- and off-target criteria. The cut-off levels were specific for each Ab and ranged from R.Z-scores of 8.7 to 195.7 (mean 20.5). Applying these Ab-specific cut-offs, we annotated each Ab for binding to the on-target, binding to the on-target and off-target(s), detecting only an off-target or not being able to detect any GPCR targets (see **Table 1B**). To classify an Ab as exhibiting on-target binding, the mean R.Z-scores obtained from the samples expressing the intended target GPCR were required to be above the cut-off. Conversely, any “off-target” datapoint value above the threshold resulted in an annotation as “cross-reactive” Ab.

**Fig. 3.**
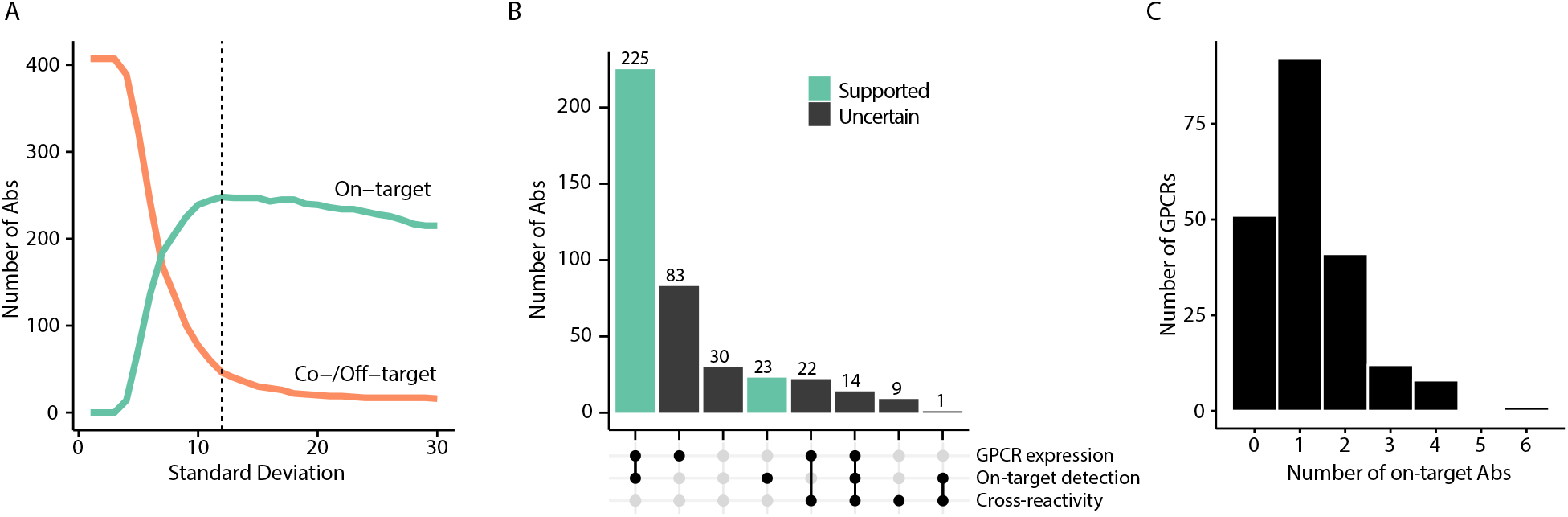
Ab selectivity threshold and summary statistics. **(A)** Data-driven selection of a hit threshold for defining Ab selectivity. The plot shows the theoretical number of Abs that would be categorized as on-target (green) and Abs that exhibit cross-reactivity (binding of unintended GPCRs, orange) as a function of standard deviations from the population peak. The threshold was selected to align with the plateau and corresponds to 12 SDs (vertical dashed line). **(B)** The numbers of Abs that fall into different categories based on evidence of GPCR expression, target detection, and cross-reactivity. Green bars indicate Abs that captured only the intended GPCR target, irrespective of the level of GPCR expression. **(C)** Histogram showing the distribution of the number of validated Abs per GPCRs.

**Table 1B.** Numbers of Abs and target specificity percentages per subfamily of target GPCRs. On-target Abs are target specific without cross-reactivity towards unintended GPCRs. Co-target Abs recognize both intended and unintended GPCRs. Off-target Abs recognize only unintended GPCRs. Abs with no target did not detect anything above the threshold.

| Subfamily | Abs [N] | On-target [%] | Co-Target [%] | Off-target [%] | No Target [%] |
| --- | --- | --- | --- | --- | --- |
| Rhodopsin (alpha) | 124 | 74.2 | 1.6 | 8.9 | 15.3 |
| Rhodopsin (beta) | 54 | 59.3 | 1.9 | 14.8 | 24.1 |
| Rhodopsin (gamma) | 98 | 54.1 | 5.1 | 4.1 | 36.7 |
| Rhodopsin (delta) | 72 | 54.2 | 2.8 | 4.2 | 38.9 |
| GSAF | 44 | 63.6 | 2.3 | 4.5 | 29.5 |
| Other | 15 | 26.7 | 26.7 | 20.0 | 26.7 |
| Total | 407 | 60.9 | 3.7 | 7.6 | 27.8 |

We found that 61% (248 of 407) of Abs tested recognized only their intended GPCR. These Abs did not exhibit any off-target enrichment above the selectivity cut-off. Out of the remaining 159 Abs, 9% (15 of 159) of Abs enriched the intended target and at least one off-target. Another 20% (31 of 159) of Abs bound only an unintended target(s), while 71% (113 of 159) did not enrich any target above the cut-off (**Fig. 3B**). The 248 highly selective Abs corresponded to 154 unique GPCR targets. Many GPCRs had two or more validated Abs and, in some cases, up to six Abs (**Fig. 3C**). Of the 407 tested Abs, we also tested 8 binders developed by sources other than the HPA. Out of these eight CABs, four Abs were highly selective, corresponding to four unique GPCR targets.

To test if any of these observations were linked to a subfamily, we determined the success rates of GPCR expression and recognition (**Table 1B, Table S2**, and **Fig. 2**). Out of the 215 GPCRs, 182 (84.7%) passed the expression threshold, and the success rate for expressing GPCRs was the highest for the rhodopsin alpha subfamily (93.2%) and the lowest for the rhodopsin delta subfamily (73.9%). Considering only those GPCRs deemed to be expressed in statistically sufficient quantities and excluding the smallest subfamily of “other GPCRs,” the chances of detecting a GPCR ranged from 71.7% to 92.5%. Successful expression of the GPCRs increased the likelihood of antigens of finding on-target Abs.

### Paired antibodies and different GPCR epitopes

One valuable approach to validate Abs is to compare several (paired) Abs that bind to different epitope regions on a common target. Among the 205 GPCRs, 116 (57%) were targeted by two or more paired Abs. This allowed us to compare binding characteristics to a GPCR, and present different scenarios for selective and non-selective recognition of solubilized receptors. Overall, 62 of 116 (53%) of GPCRs with paired Abs were recognized by ≥ 2 Abs. Agreement between paired Abs was evaluated using Pearson correlation between pairs and was found to be elevated (mean of r = 0.71; median of r = 0.89).

To exemplify a few scenarios of paired Abs, we selected GPCRs with the highest numbers of paired Abs per respective family. We summarized the performance of each Ab in recognizing solubilized receptors in **Fig. 4** and **fig. S2**. For the rhodopsin alpha subfamily, adrenoceptor alpha 2B (ADRA2B) was selectively recognized by six out of six Abs. They were raised against three distinct epitope regions, all within the intracellular loop (ICL)3 of ADRA2B. For the rhodopsin beta subfamily, gastrin-releasing peptide receptor (GRPR) was selectively recognized by five Abs out of six. The Abs were raised against two distinct epitope regions, one within the extracellular domain (ECD) (HPA059693, HPA069267, and HPA077564), and one within the extracellular loop (ECL)2 (HPA059022, HPA069604, and HPA077557). HPA059693 did not capture the intended target, while the other Abs raised against the same ECD-based antigen, HPA069267 and HPA077564, were on-target only. For the delta subfamily, C-C motif chemokine receptor 7 (CCR7) was selectively recognized by only one out of five Abs even though three distinct antigens were used: one antigen in the ECD, one antigen in ECL2 and one corresponding to the intracellular C-terminal tail of the receptor. HPA060045 was the only Ab tested with selective recognition of CCR7. The other two Abs raised against the same ECD antigen did not detect any target above the threshold. The expression levels of CCR7 were lower than most receptors of the delta subfamily and hence classified as “uncertain”. This could indicate that only HPA060045 has a sufficiently high affinity to CCR7 to detect even lower levels of this GPCR. For the rhodopsin gamma subfamily, follicle-stimulating hormone receptor (FSHR) was selectively recognized by three out of five Abs with less distinct Z-score differences than in the previous examples. HPA067689 and HPA078372 were raised against a common antigen close to the first transmembrane helix (TM1) of the N-terminal tail and failed to capture FSHR. The on-target Abs were raised against two antigens further towards the N-terminus of the ECD. The three distinct epitope regions are all within the large ECD of FSHR. For the glutamate subfamily, GPCR class C group 5 member D (GPRC5D) was selectively recognized by three out of four Abs. The failed Ab HPA047203 was raised to the ECL2 region, while the selective Abs targeted its ECD (HPA071909) or intracellular C-terminal tail (HPA064241 and HPA071739).

**Fig. 4.**
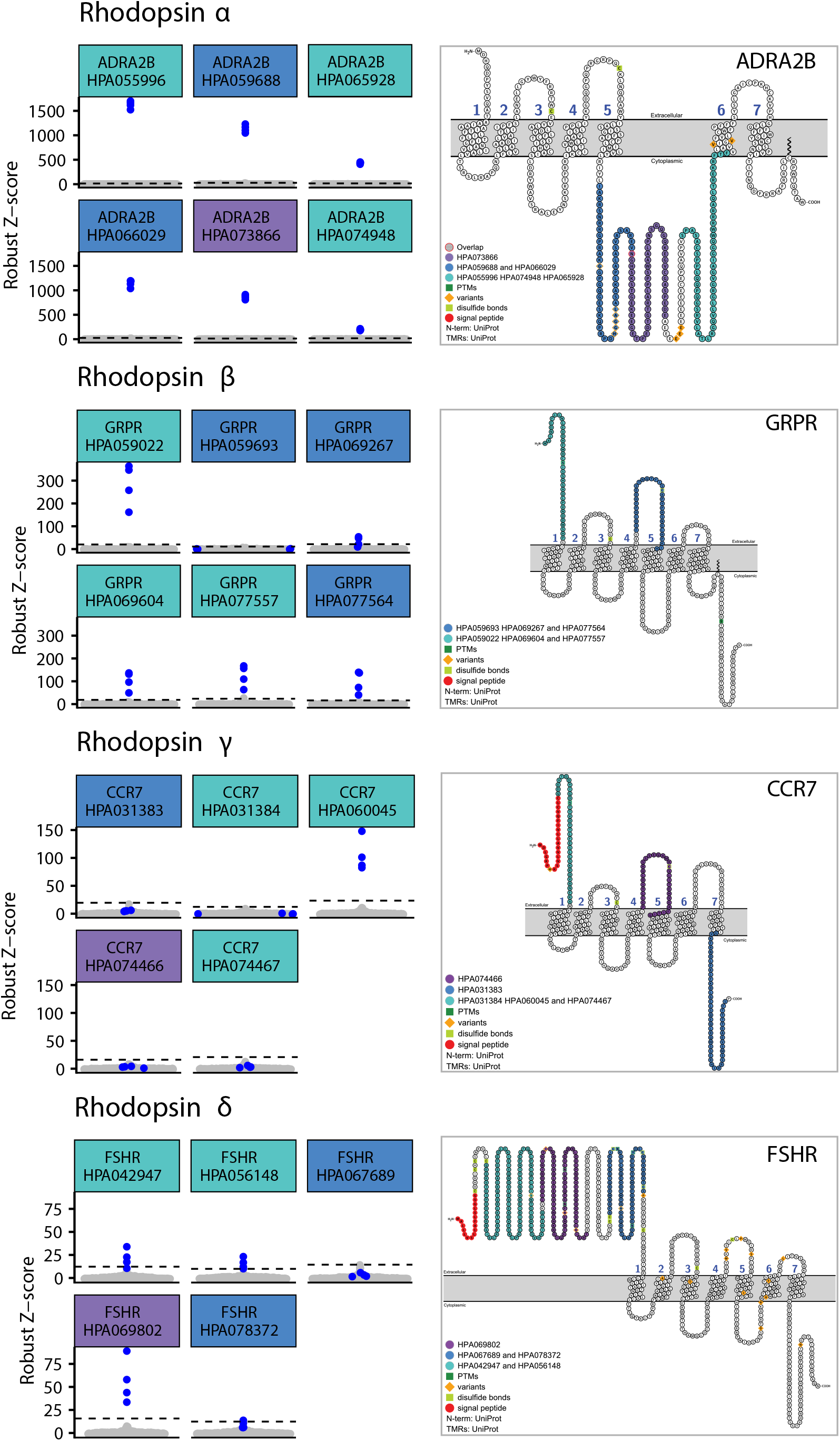
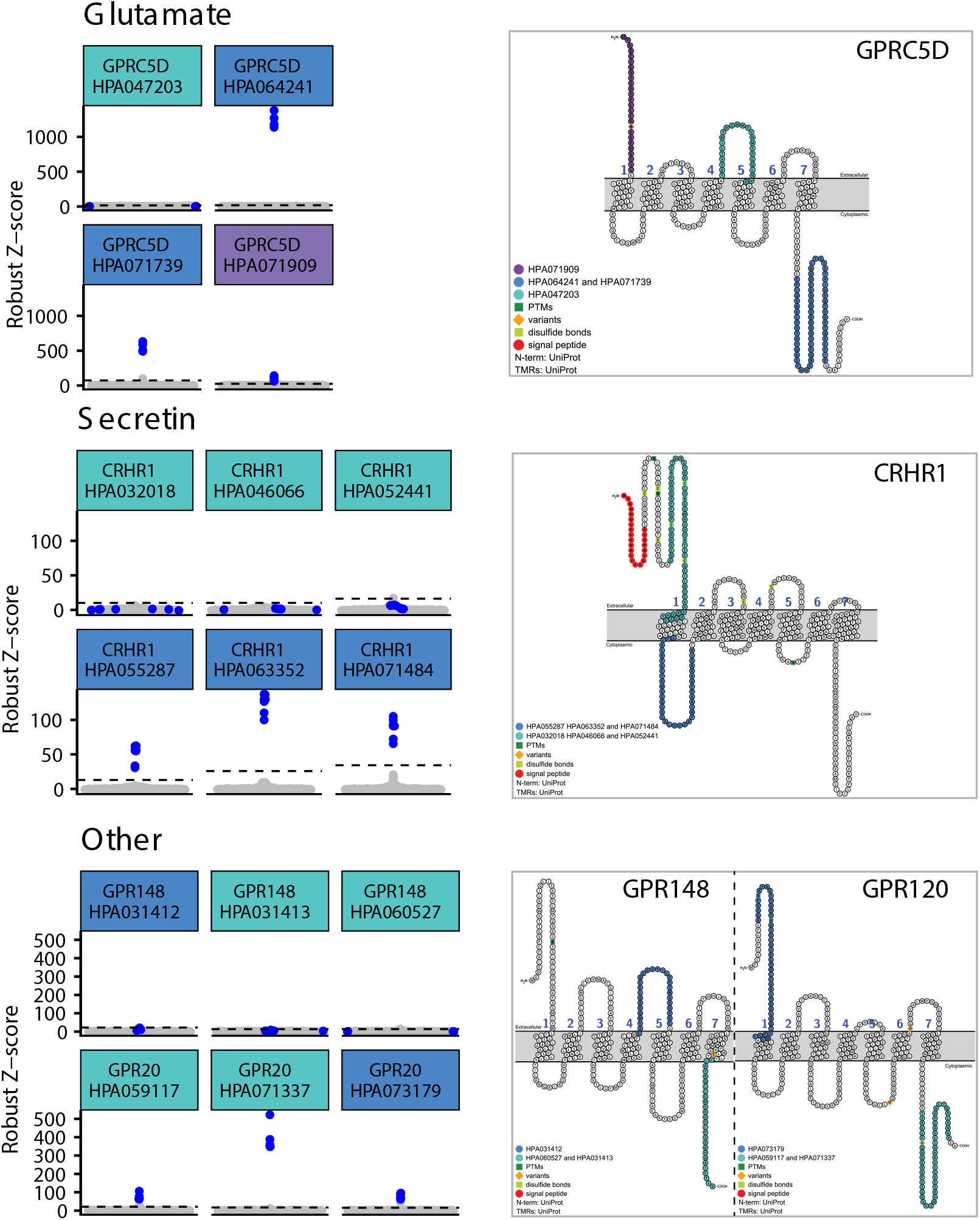
Detection of GPCRs with paired Abs. Left column: Beeswarm plots showing binding events for multiple Abs targeting the same GPCR. Intended GPCR, blue dots; unintended GPCRs, grey dots. Dashed lines correspond to the selectivity cutoff for each HPA Ab. Color coding of HPA Ab ID corresponds to the color coding of the antigen in the snake plot diagram. Right column: Snake plot diagrams showing the antigen sequence used to generate the antibody on the entire protein sequence. Some Abs have the same antigen sequence. Generated with Protter (*36*). Both columns are divided per subfamily.

We found cross-reactivity of HPA071739 for GPRC5A, albeit at a relatively low Z-score compared with the Z-scores for GPRC5D-containing samples. The Ab raised against the same antigen, HPA064241, was highly selective for GPRC5D only. For the secretin-like subfamily, three out of six Abs tested for corticotropin-releasing hormone receptor 1 (CRHR1) were on target. Selective recognition of CRHR1 was linked to an antigen representing the ICL1. Surprisingly, the three HPA Abs generated against a 60-residue-long antigen on the ECD did not generate Abs to detect solubilized CRHR1 in the assays. For the FZD subfamily (**fig. S2**), Abs against frizzled receptors 4 and 5 (FZD4 and FZD5) provide examples of differential Ab recognition. Here, one of two tested Abs detected the target proteins, respectively. For FZD4, the selective Ab was raised against a shorter intracellular C-terminal tail antigen (HPA042328) rather than a longer N-terminal one corresponding to the ECD (HPA074833). For FZD5, the Ab targeting the shorter and slightly more N-terminal sequence within the ECD was the selective one (HPA052361), although the other Ab (HPA053811) was also raised against an ECD antigen.

Lastly, GPCRs grouped as “other” are represented by GPCR 20 (GPR20) and GPCR 148 (GPR148). For GPR20, all three Abs captured the solubilized receptor, but we observed crossreactive binding of HPA059117 to the arginine vasopressin receptor 1B (AVPR2) and of HPA073179 to the PRLHR receptor. Interestingly, HPA059117 and the selective Ab HPA071337 were raised against the same intracellular C-terminal tail antigen, while the antigen of HPA073179 was within the ECD and extended slightly into TM1. For the receptor GPR148, none of the three tested Abs recognized the solubilized or any other target above the set Z-score threshold. The antigens for the Abs are located within the ECL2 and intracellular C-terminal regions of GPR148.

### Deconvolution off-target binding

A subset of 46 out of 407 Abs recognized GPCRs other than the intended targets. To investigate possible reasons for their unspecific performance, we checked the sequence homologies and differences in expression levels between the on- and off-target GPCR. As shown in **Fig. 5**, differences in GPCR abundance and sequence similarity contributed to the off-target binding. For approximately half of the Abs, a twofold higher relative abundance of the off-target led to its recognition. For specific cases, lower off-target abundance was compensated by high sequence similarity (E-value < 1) (**Fig. 5A**). In other cases, we also observed a slight difference in off-target GPCR expression within a subfamily (**Fig. S3**). Overall, there was a similar percentage of Ab cross-reactivity (10-20%) for all subfamilies except “other” (**Fig. 5B**). The latter group may have a higher percentage of off-targets because (i) it was also the smallest group, (ii) the receptors in this group are of unknown phylogenetic positioning, and (iii) uniquely, we tested these Abs against all GPCRs in the library.

**Fig. 5.**
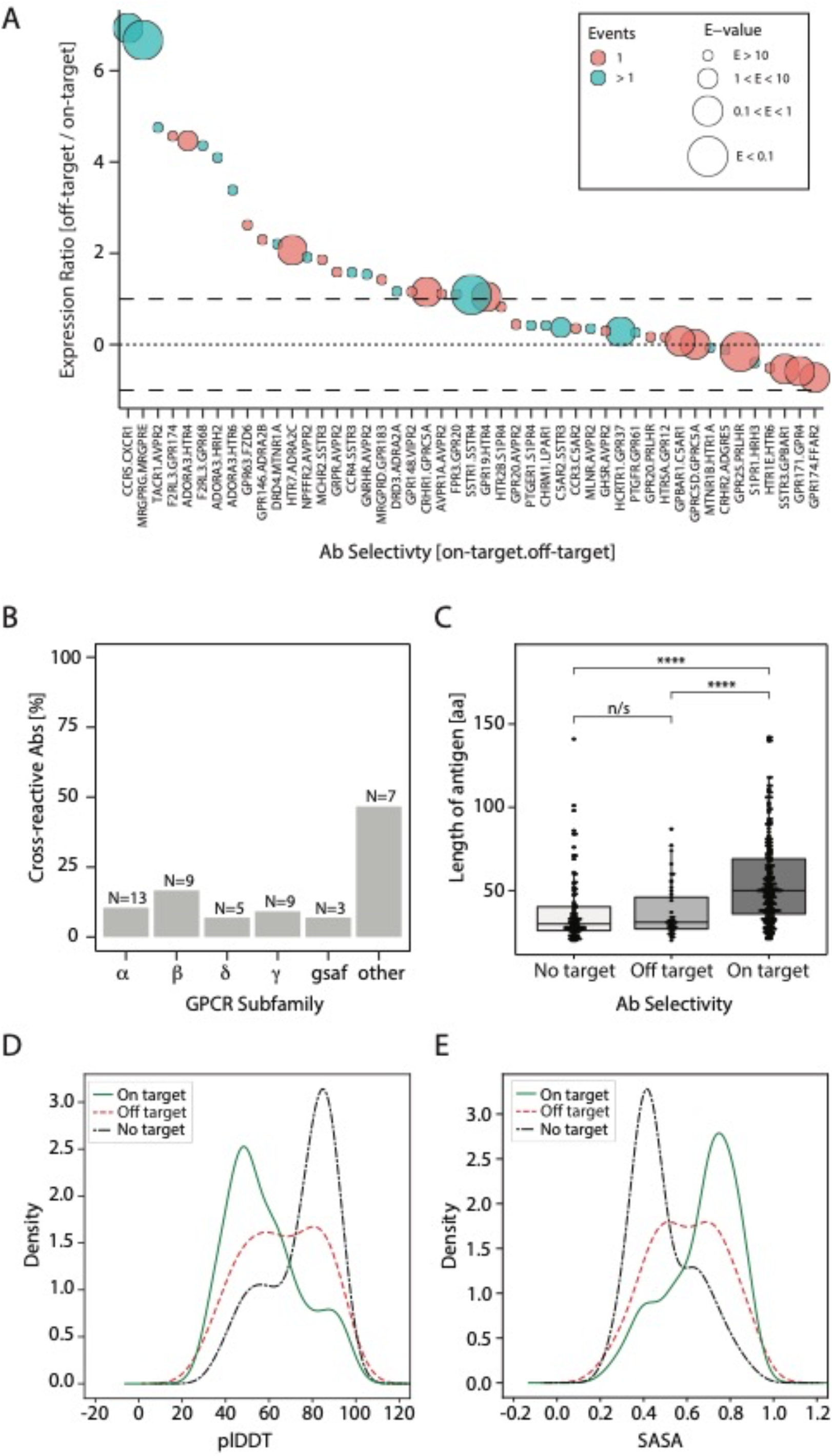
Deconvolution of observed Ab cross-reactivity behavior. **(A)** Deconvoluting off-target detection through analysis of expression ratio and sequence homology. The expression ratio between each on-target GPCR (denominator) and off-target GPCR (numerator) is plotted from highest to lowest. The x-axis represents Abs binding off-target and displayed by their “on-target.off-target” GPCR. The black dashed line indicates a two-fold difference in expression ratio. The dotted line corresponds to an equal on-target and off-target GPCR expression. The size of each dot conveys the E-value of the particular target GPCR and off-target GPCR pair. The lower the E-value, the bigger the circle and the more similar the GPCRs are to each other in the primary sequence. If the off-target GPCR was only captured in one sample, the dot is peach; if the off-target GPCR was captured in more than one sample containing it, the dot is colored teal. **(B)** Summary of the number of cross-reacted GPCRs per group tested, shown as a percentage of total Abs per group and as an absolute number. **(C)** Comparison of the antigen length used for producing the HPA Abs binding on-target (N=245) versus those annotated to bind co/off-target (N=45) or binding no-target (N=109). A two-sided Wilcoxon rank sum test determined significant differences between the three groups (p=2.5 × 10^−15^). Here, **** indicates a p < 1 × 10^−6^; n/s refers to “not significant”. **(D)** The predicted local Distance Difference Test (plDDT) was conducted for the three Ab selectivity classes to determine confidence in structures predicted for the respective antigens on full-length proteins. The average plDDTs of antigens from on-target Abs (green line) are predominantly < 50 and indicated disordered structures. Predictions for antigens from off-target Abs are shown in a dashed red line and those from no-target Abs in a black and dot-dashed line. **(E)** The averages of solvent-accessible surface areas (SASA) were calculated for antigens from the three Ab selectivity classes to determine the accessibility of the antigens on the full-length proteins. Based on the ranks, antigens from on-target Abs (green line) are more exposed than off- or no-target Abs. Average SASA values for antigens from off-target Abs are shown as dashed red and no-target in black and dot-dashed lines.

Next, we investigated whether any systematic pattern in off-target recognition could be observed. As shown for examples in **Fig. S4**, ten Abs exhibited consistent off-target binding in all replicated samples expressing the off-target GPCR. We observed such examples in five subfamilies. The Ab for C-C motif chemokine receptor 5 (CCR5) detected the C-X-C motif chemokine receptor 1 (CXCR1) in all samples. Identifying these off-target binders provides further support and validity for our approach.

The cross-reactivity of most Abs could be attributed to sequence homology or large differences in expression ratio. The third category of Abs, however, cross-reacted neither due to abundance nor the GPCR sequence homology. We deemed these unselective Abs promiscuous and generally unsuitable for the GPCR analysis in the presented application. We also tested other criteria that might influence whether an antibody bound on-target or not. Considering that all HPA Abs are processed via one pipeline, we checked whether the length of the antigen used to generate the Abs influenced the success rate (**Fig. 5C**). We found a significant difference between the antigens used to generate on-target Abs (*P* = 1.1x 10^−13^). These antigens were generally longer (54 aa ± 26; *N* = 248) compared with those that generated Abs that bound off-target or that did not recognize any GPCR (37 aa ±19;*N* = 159).

We analyzed the predicted structures from AlphaFoldDB (*22, 23*) of all the antigens to analyze the properties of the epitopes causing Abs to be specific binding or not (**Table S3**). On-target binders are significantly (P=1.7×10^−14^) enriched in protein regions predicted to be disordered (as defined by a low plDDT score) (**Fig 5E**). Further, the on-target epitopes are also enhanced in residues exposed to the surrounding (P=1.1×10^−15^) **Fig 5F**, enriched in coils (P=3.2×10^−9^), and depleted in helices (P=6.6×10^−5^) and sheets (P=6.2×10^−6^).

### Interactive selectivity analysis

To enable better access to our data, we developed a web-based interface (**Fig. 6**). The app can be accessed at [URL: *provided upon publication*] and allows interactive Ab- and GPCR-centric browsing of the assay results and includes information about GPCR expression and enrichment of GPCR per Ab. It presents a summary of the selectivity analysis and allows users to display heatmaps to overview the Ab selectivity per GPCR subfamily. The app also shows correlation analyses of paired Abs raised against a common GPCR.

**Fig. 6.**
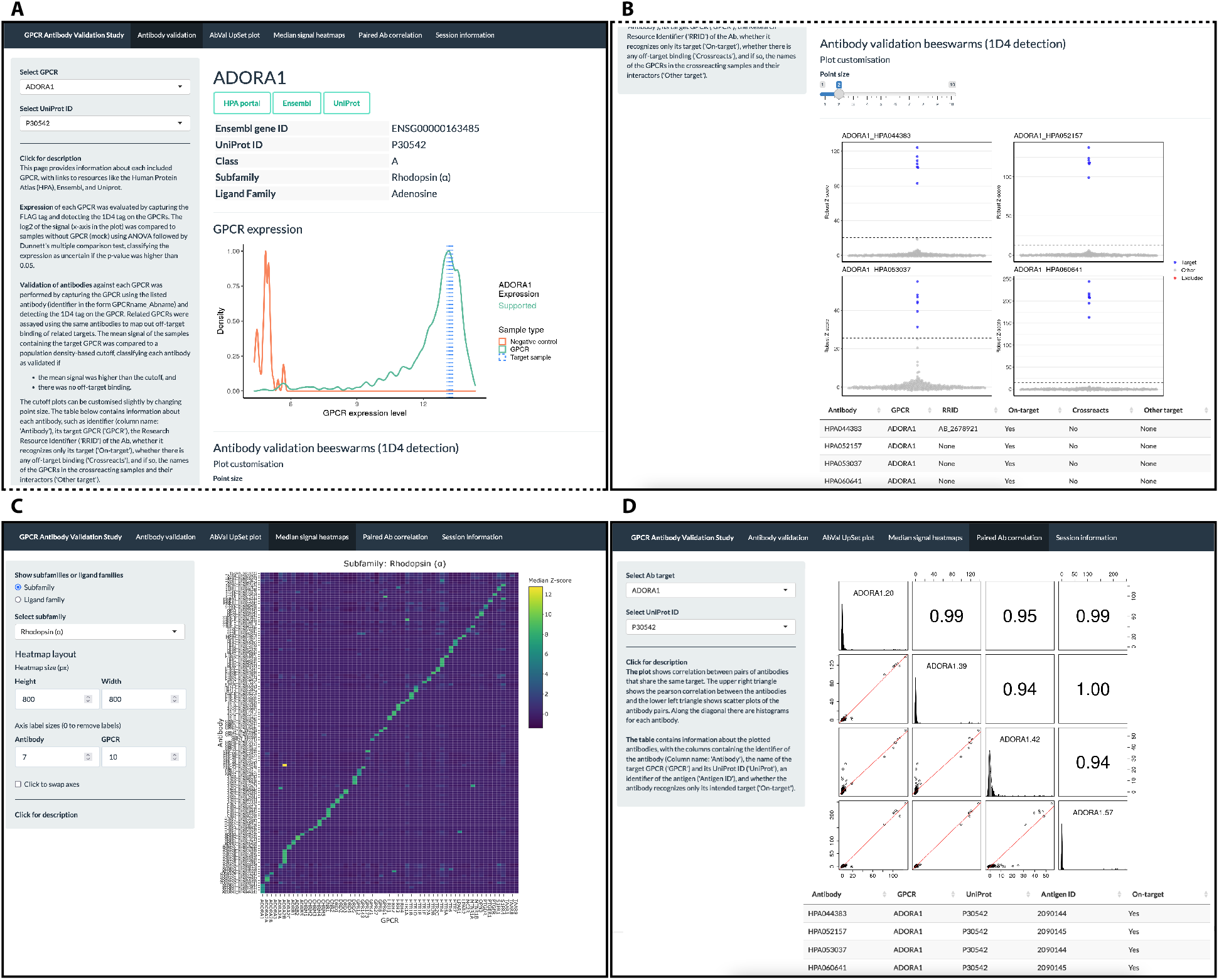
A brief overview of the web-based interface. **(A-D)** The interactive web-based interface contains tabs with information about each Ab. **(A)** The expression of each GPCR visualized as expression density plots and (**B**) the performance of Abs targeting it, visualized as beeswarm plots. **(C)** Cross-reactivity of Abs with phylogenetically related targets visualized as heatmaps. **(D)** Comparison of paired Abs. The interface was generated through the *shiny* R package.

## DISCUSSION

Most of the Abs used in our study were generated by the HPA project that used a bioinformatic algorithm to select unique features of the primary protein structures to produce highly target-specific antigens (*16*). Newer computational tools have the potential to generate further improvements in antigen designs. The scope of machine learning applications has expanded dramatically within the past three years and accelerated the prediction of protein structures. The advent of Alpha-Fold (*22*) has recently been harnessed by others to create *in silico* models for designing binding reagents for any interface (*24*). It will be interesting to follow the development of such approaches and compare the experimental validation data accompanying computational classification schemes. For the time being, mapping the epitope regions of paired Abs onto two-dimensional GPCR snake diagrams can be complemented by mapping the access to the epitopes in predicted 3D structures. Our analysis highlights that it is unfavorable to generate antibodies against regions of antigens that are buried. For example, based on the solved structure of the ECD of CRHR1, it appears that a portion of the antigens corresponding to the three failed anti-CRHR1 Abs (HPA032018, HPA046066, and HPA052441) folds into two beta-sheets (*25*). The distinct structural features on the GPCR may prevent the Abs raised against antigens mapping to that structural region from recognizing it. Interestingly, the most successful antigens appear to be disordered regions, providing a direction to be further explored. Nonetheless, the structural prediction of GPCRs and other membrane-bound proteins remains challenging compared to soluble proteins.

A useful feature of our approach was to include expression levels into our assessment scheme. Looking at the GPCR expression levels for the different SBAs, rhodopsin alpha appears to have the highest percentage of expressed GPCR and is also the biggest (N=59, 93.2% expressed). In comparison, rhodopsin delta and “other” groups had the lowest overall expression (73.9% and 66.7%). The “other” GPCR grouping contained only nine receptors, so it may not be a conclusive comparison. Rhodopsin delta subfamily GPCRs, of which many are orphan receptors, had an overall lower success rate in expression compared to the alpha receptors. In our library, 47.8% of the rhodopsin delta subfamily receptors are classified as orphans. In comparison, 13.6% of rhodopsin alpha subfamily receptors, 4.8% of rhodopsin beta subfamily receptors, 3.9% of rhodopsin gamma receptors and 20.7% of GSAF receptors are annotated as orphan. Unsurprisingly, 100% of the “other” GPCRs are orphans. Alternative expression schemes offer possible routes to improve expression levels. Still, the low expression yields of orphan receptors can be seen as one of the reasons why endogenous ligands for some of the GPCRs have not yet been identified.

Multiplexed planar protein arrays were used previously to test Ab selectivity (*18*). Recent efforts by Syu et al. have also resulted in similar arrays for GPCRs (*26*). There are, however, key conceptual differences between the planar and suspension bead arrays: Are the GPCRs or the Abs immobilized or added in solution? It is worth noting that immobilizing and drying the target may alter its molecular integrity and limit the accessibility of some epitopes, in particular when maintaining sensitive or embedded structures. In the SBA, micelle-imbedded GPCRs are generated from detergent solubilization, which is highly representative of GPCRs found in cellular membranes. From an analytical perspective, planar and bead arrays follow the ambient analyte assay theory (*27*). For planar arrays, the Abs bind and rebind on the area where a target is immobilized. For the SBAs, the Abs capture targets that can rebind on the Ab-coupled surface. While Abs are regarded as molecules with a rigid structure, membrane-imbedded GPCRs are known to be structurally labile. Our data suggest that preparing and maintaining the integrity of GPCRs, both as on- or probable off-target, is an essential benefit of the SBA approach. It also enables modulating the GPCR with a higher degree of freedom. Capturing the GPCRs with epitope tags also allowed us to ensure that sufficient target molecules were present in each assay. This approach also allows for the future use of such GPCRs as targets to assay for the presence circulating auto-Abs in serum samples (*28, 29*).

We built our approach on polyepitope pAbs that were readily available for many applications from the HPA project(*30*). The sustainability of pAbs is low due to the limited volumes obtained from the Ab generation process. Still, there are examples where functional mAbs have been produced on antigens identified from pAbs (*31*). The polyepitope characteristic of pAbs may have also contributed to the high success rate observed, as one epitope can compensate for the inaccessibility of another one. The HPA Abs are generated against protein fragments prepared and stored in high-content urea. The fragment length and storage conditions may limit the antigen’s ability to form delicate tertiary structures and represent the native antigen region more closely. Nonetheless, many HPA Abs were found to recognize the overexpressed GCPRs in micelles, so we wanted to check how well these Abs bound to the endogenous receptors on fixated and denatured tissues or cells. Of the 400 anti-GPCR Abs investigated here, 97 Abs (24%) have been published on the HPA portal (version 21.1). These Abs have been classified as supported or approved by the stringent enhanced validation criteria that focus on the Abs’ utility to stain tissues and cells: 28% (73/261) have passed the test for immunohistochemistry (IHC) (*9*), 12% (46/399) for confocal microscopy (*10, 11*) and 22% (83/370) of the Abs were classified as supportive for the use in Western blot (URL: https://v21.proteinatlas.org/about/antibody+validation). In one example, we observed good agreement between the results of the SBA assay and HPA validation by immunofluorescence and Western blot for an Ab targeting the sphingosine-1-phosphate receptor 4 (S1PR4) (**fig. S5**). In general, the differences in the utility of the Abs confirm the influence of assay- and sample-specific conditions and support the need to apply appropriate validation schemes.

The reasons why we observe a 60% success rate in validating the Abs can be explained by a combination of factors. Importantly, all the mentioned aspects apply equally to the testes in which we incubated off-target GPCRs with the Abs. Over-expression of the target reduces the sensitivity burden, as the on-target becomes the dominant protein, and interfering proteins may not be able to occupy the GPCRs. Preparing the samples as membrane fractions reduces their complexity and lower the selectivity burden for the Abs. In addition, removing non-membranous but abundant cellular proteins reduces the probability of binding to other non-GPCR off-targets. Using detergent to solubilize the receptors preserves epitope accessibility to facilitate in-solution immunocapture of the GPCRs and circumvents the need to fixate, cross-link, or dry GPCR-containing membranes. In addition, we attribute rebinding events on the Ab-coupled beads to the validation outcome. Consequently, the high rate of target selectivity suggests that the chosen assay conditions support Ab recognition of their intended target among several over-expressed potential off-targets.

Our method of measuring GPCR expression across all samples relies on selective Ab recognition of the FLAG and 1D4 epitope tags engineered into each receptor. FLAG tyrosine sulfation was previously reported to affect GPCR expression and, if present, may result in a “false negative” determination of GPCR expression (*32*). Although the work by Hunter and colleagues focused on a dopamine receptor, and all five dopamine receptors in our library showed good expression, it is possible that this phenomenon affected other GPCRs that expressed more poorly. Abs validated here are not necessarily specific in other applications, such as those in which solubilization is not possible or where the GPCR epitopes are presented in a different state. We used overexpression to present sufficient quantities of the intended target, as well as all other targets, to the surface-bound Abs. Ectopic expression changes the proportion of target abundance over all other proteins in the micelles. It also presents the GPCRs to the Abs at levels that are likely non-physiological. Thus far, we have mostly tested HPA Abs, but the approach is not limited to these and can be expanded to other Abs, nanobodies, or scaffold proteins if these can be immobilized to the beads. Caution must be raised when smaller molecules are used because coupling these to a solid support may limit their functionality. Our approach was built on fulllength GPCRs embedded into detergent-lipid micelles. Hence it does not give us the resolution to determine the exact binding epitopes down to the amino acid level. As of today, we cannot predict a success rate for reproducing selective pAb and if these will be functional in the assay even if the antigen it was raised against has previously generated a validated pAb. Hence, the discovery and validation of newly developed Abs will still require molecular tests. Preferably the analytical protocols and assay conditions for the most appropriate applications have already been defined.

## MATERIALS AND METHODS

### Experimental Design

The objective of this study was to determine the selectivity of anti-GPCR Abs. Each binder was tested in an SBA assay against detergent-solubilized and overexpressed GPCR from Expi293F cells. The selectivity of the Abs was tested for their intended GPCR and against GPCRs from the same subfamily. Abs were coupled to color-coded beads, and their binding to a GCPR protein was detected via fluorescently labeled Abs specific for epitope tags. Per Ab-coupled bead ID, the median fluorescence intensity (MFI) of at least 32 events was used as data for the selectivity analysis.

### Materials

Information on all Abs used in the SBA generation can be found in **Table S1**. Abs were either from HPA (some of which are commercially available from Atlas Antibodies AB) or purchased from Affinity Biosciences. Expi293F cells were a gift from the Ravetch Lab (The Rockefeller University. Expi293 Expression Medium (cat. A1435101) and the Expifectamine 293 transfection kit (cat. A14524) were from Fisher. The cells were cultured in 125-mL flasks (cat. 431143), 250-mL flasks (cat. 431144), and 12-well plates (cat. 353043) from Corning. PE-conjugated anti-FLAG Ab was from BioLegend. Anti-1D4 Ab and anti-OLLAS Ab were conjugated to PE using an Ab conjugation kit from Abcam (cat. 102918) according to the manufacturer’s protocols. Half-area 384-well plates were from Greiner. Blocking reagent for ELISA (BRE, cat. 11112589001) was from Roche. DC Assay kit was from Bio-Rad. Phosphate-buffered saline (PBS) was from Medicago. ProClin 300 (cat. 48912-U), cOmplete mini protease inhibitor tablets (cat. 11836170001), casein (cat. C7078), polyvinyl alcohol (PVA, cat. 25213-24-5), polyvinylpyrrolidone (PVP, cat. 9003-39-8), and FLAG M2 Ab (cat. F3165) were from Sigma-Aldrich. n-dodecyl-b-D-maltoside (DM, Cat. D310, CAS 69227-93-6) was from Anatrace. Purified rabbit IgG was from Bethyl (P120-101). Anti-mouse IgG- and anti-rabbit IgG-conjugated R-PE were from Jackson ImmunoResearch (cat. 115-116-146 and 111-116-144, respectively).

### Cell Culture and Transfection

Expi293F were cultured and transfected according to manufacturer’s instructions. Briefly, cells were cultured in serum-free Expi293 Medium using culture flasks under constant shaking at 130 rpm at 37°C with 8% CO2. For transfection, cells were counted using a Nexcelom Cellometer Auto T4 and diluted to 2,000,000 cells/mL and were allowed to grow overnight. The next day, the cells were counted, diluted to 3,000,000 cells/mL, and 1.25 mL of cells was transferred to each well of a 12-well culture plate. Transient transfections were then performed with the Expifectamine 293 transfection kit (Thermo Fischer Scientific). Each well of cells was transfected with 4 μl of FreeStyle MAX Reagent and 0.25 μg of GPCR plasmid DNA. Total transfected plasmid DNA was kept constant at 1.5 μg/well by adding empty vector pcDNA3.1(+). Enhancers were added 18-24 hours after transfection according to the manufacturer’s protocol and cells were harvested 72 hours after transfection.

### DNA constructs

Epitope-tagged human GPCR DNA constructs were encoded in a pcDNA3.1(+) mammalian expression vector. All GPCRs except CALCRL and FZD4,5,6, and ten relied on the codon-optimized PRESTO-tango library of signal sequence-FLAG-GPCRs as a starting point to generate 215 FLAG-GPCR-1D4 constructs, with the FLAG tag (DYKDDDDA) following the HA (hemagglutinin) signal sequence MKTIIALSYIFCLVFA. C-terminal components of the original PRESTO-Tango constructs were removed (V2 tail, TEV site, Tta transcription factor) upon addition of 1D4. The amino acid sequence of the C-terminal 1D4 tag is DEASTTVSKTETSQVAPA. The PRESTO-Tango plasmid kit was a gift from Bryan Roth (Addgene kit # 1000000068).

All GPCRs retained their endogenous signal sequences, if applicable, in the original PRESTO-Tango library, thereby having two signal sequences in the final plasmid. We removed the endogenous signal sequence for 13 receptors (F2R, GABBR1, GLP1R, GPR156, GPR37, GPR97, GRM1, GRM2, GRM4, GRM5, GRM6, GRM7), but not from 11 receptors (CALCR, CD97, CRHR1, CRHR2, GCGR, GIPR, GPR114, GPRC5B, GPRC5C, GPRC6A, VIPR2).

There were no frizzled receptors in the PRESTO-tango library, so we included four in-house frizzled receptors. The human frizzled GPCRs FZD4, FZD5, FZD6, and FZD10 cDNAs encode the 23–amino acid residue 5-hydroxytryptamine receptor 3a receptor (5-HT3a) signal sequence (MALCIPQVLLALFLSMLTGPGEG) in place of the native signal sequence. There is a modified amino acid on position 2 (A instead of R) to optimize the Kozak sequence to GCCGCCACCATGG. An HA tag follows the signal sequence. C-terminal to the receptor is a 1D4 tag. The cDNA for the GPCRs FZD4, 5, 6 and 10 were designed in-house and synthesized through Genewiz.

### Clarified lysate preparation

Cell membranes were solubilized as previously described (*15*). Briefly, cells were solubilized with DM detergent to form micelles around membrane proteins and maintain GPCR structure. 72 hours after transfection, Expi293F cells were harvested and washed twice with cold PBS. Cells were then incubated in solubilization buffer [50 mM HEPES, 1 mM EDTA, 150 mM NaCl, and 5 mM MgCl2 (pH 7.4)] with 1% (w/v) DM and cOmplete mini protease inhibitor for 2 hours at 4°C with nutation. Following solubilization, lysates were clarified by centrifugation at 22,000g for 20 min at 4°C. Solubilized lysates were then transferred to a microcentrifuge tube, and total protein content was determined by Protein DC assay according to the manufacturer’s specifications. Solubilized lysates were flash-frozen prior to storage.

### Suspension Bead Arrays

Anti-GPCR and anti-tag Abs were covalently coupled to color-coded magnetic beads (MagPlex, Luminex Corp.) as previously described (*15*). In short, 1.75 μg of each Ab was diluted in MES buffer [100 mM 2-(*N*-morpholino)ethanesulfonic acid (pH 5.0)] to a final volume of 100μL. The diluted Abs were then conjugated onto the carboxylated beads using NHS-EDC chemistry. After washing away unbound antibodies, the reactions were quenched with BRE buffer overnight. The Ab-coupled beads were subsequently grouped and pooled to form six subfamily-related SBAs. The majority of the Abs were rabbit pAbs, and the coupling efficiency was determined using anti-rabbit-RPE Abs (Jackson ImmunoResearch) and the data was collected using a FlexMap 3D instrument (Luminex Corp., xPONENT Software, build 4.3.309.1).

### Assay procedure

The procedure described here is based on and consistent with that of our proof-of-concept study (*15*); Clarified protein lysate were diluted to 2 μg/μl in solubilization buffer (described above) with 0.01% (w/v) DM in a 96-well plate, and diluted again 3.6 times in SBA buffer such that 12.5μL of lysate was combined with 32.5μL of buffer (PBS containing 0.5% PVA (w/v), 0.8% PVP (w/v), 0.1% (w/v) casein, and 10% rabbit IgG). We then transferred 45μL of the solution to a 384-well assay plate containing 5 μl of bead array using CyBio SELMA (Analytik Jena). The lysates and beads were incubated overnight (16 hours) at 4°C. Next, the plates were washed six times with 60 μl of PBS containing 0.05% Tween 20 (PBST) using a BioTek EL406 washer. Detection was enabled by the addition of 50μL PE-conjugated anti-tag Abs diluted in BRE containing 0.1% DM, 0.1% Tween 20, and 10% rabbit IgG and the plate incubated for 1 hour at 4°C. The final dilution used for the detection Ab (PE-conjugated anti-1D4) was 1:1000. The beads were washed six times with 60μL of PBST. After the final wash, 60μL of PBST was added to the beads, and the fluorescence associated with each bead was measured using a Luminex FlexMap 3D. The data are reported as MFI.

### Web-based interface

A web-based companion R app to the study was made using the *shiny* package (version 1.7.1) and R version 4.2.0, containerized using *Docker* (version 20.10.16, build aa7e414) and hosted on the SciLifeLab Serve platform. The *Shiny* app was created to contain information about each assayed GPCR, such as GPCR expression and on-target and off-target Abs for each, an overview of the validation status of different Abs, interactive heatmaps made using the *plotly* package (version 4.10.0) showing the reactivity of each Ab against the GPCRs they were used against, and correlations between pairs of Abs that share the same target using the *paired.panels* function of the *psych* package (version 2.2.5). All packages and versions used for visualization are listed within the app.

### Statistical analyses

All data analysis was performed using R version 3.6.0 and plots were produced using the *ggplot2* package (version 3.3.6), unless otherwise stated. GPCR expression was evaluated in using MFI data from FLAG capture and 1D4 detection with significance testing by ordinary one-way ANOVA (*aov* function of the *stats* package, version 3.6.0) followed by Dunnett’s multiple comparison test to mock (*DunnettTest* function of the *DescTools* package, version 0.99.43). Expression per subfamily was tested first, followed by expression of each individual GPCR. GPCRs with a p-value below 0.05 were classified as having their expression supported, while other GPCRs were classified as having uncertain expression. Reproducibility of select GPCRs (ADRB2, GPRC5A, HCRTR2) was tested in biological triplicates and technical duplicates.

To bring measurements of different GPCRs to a similar scale and identify those Abs that detect only their intended target via detecting the outliers, MFI values were converted to robust Z-scores (R.Z-score) using the formula (x - median(x) / (1.4826 * MAD(x)), where × is a vector of measurements from one binder, calculated separately per binder.

The selectivity cutoff for each anti-GPCR antibody was determined by adding 12 standard deviations of the expected negative proportion of the population around the Gaussian smoothing population density peak as previously described (*33*). An Ab was classified as on-target if the mean R.Z-score of samples containing the target GPCR was higher than the cutoff value and if there were no samples with other GPCRs above the cutoff. R.Z-scores were visualized in beeswarm plots, made using the *ggbeeswarm* package (version 0.6.0). An overview of the validation was plotted in an UpSet plot showing the distribution of antibodies fulfilling different criteria. The plot was made using the *ComplexUpset* package (version 1.3.3).

Amino acid sequence similarity between antigen sequences and cross-captured proteins were compared using the blast function from *rBLAST* package (version 0.99.2) and was reported as Evalues. Alluvial plots were prepared using *ggalluvial* package (version 0.12.3). The difference in antigen lengths between on-target and off-target Abs was compared using a twotailed Wilcoxon test.

### Structural mapping

All models of the antigens were downloaded from AlphaFoldDB (*23*) using the corresponding UniProt IDs. The secondary structure and exposed surface area were extracted using DSSP (*34*). The disorder of a region was extracted from the plDDT values as this is a good indication of disorder (*35*). Statistical differences between the groups were calculated using the independent Students T-test.

## ACKNOWLEDGMENTS

We thank the Human Protein Atlas team, the Affinity Proteomics Unit at SciLife Laboratory and D. Lewis for the discussion concerning data analysis. We thank Thomas Huber for design of some of the genetic constructs used in this study.

## AUTHOR CONTRIBUTIONS

Conceptualization: IBK, TPS, JMS; Methodology: IBK, TPS, JMS; Investigation: IBK, AB; Software: LD, SF; Formal analysis: IBK, AB, LD, SF, AE, TPS, JMS; Visualization: LD, IBK, AB, JMS; Supervision: TPS, JMS; Resources: MU, TPS, JMS; Writing–original draft: IBK, AB, LD, TPS, JMS; Writing–review & editing: IBK, AB, LD, TDC, MU, TPS, JMS

## DATA AND MATERIALS AVAILABILITY

All data are available in the main text or the supplementary materials and are presented in the Shiny App and provided with this paper [URL: *provided upon publication*]. Upon publication, data and special code can be accessed on FigShare [URL: https://scilifelab.figshare.com] under the specific ID [URL: *provided upon publication*]. Additional data related to this paper may be requested from the authors. Abs used in this study are commercially available from different sources through their stated IDs. Other requests should be made to the corresponding authors.

## SUPPLEMENTARY MATERIALS

**Fig. S1.**
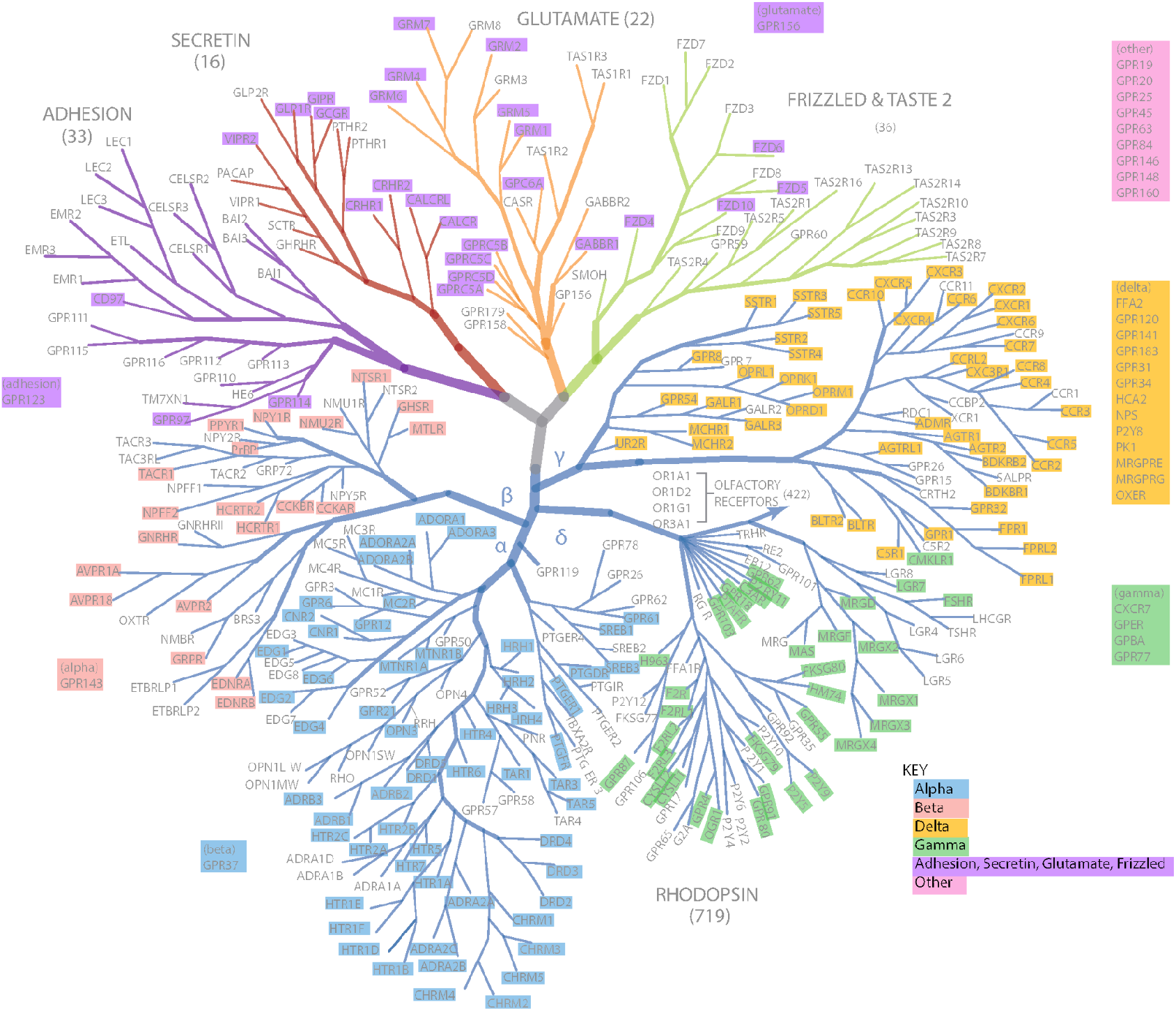
Positions of selected GPCRs on the phylogenetic tree. GPCR phylogenetic tree (*2, 37*) highlighting the 215 receptors used in this study. The color of the highlighting indicates the grouping of the corresponding Abs for each subfamily SBA. Blue, Rhodopsin family, alpha; Peach, Rhodopsin family, beta. Green, Rhodopsin family, gamma; Gold, Rhodopsin family, delta; Purple, Glutamate, Adhesion, Secretin, and Frizzed (GSAF) families; Pink, other. Adapted from Lv et al. (*38*).

**Fig. S2.**
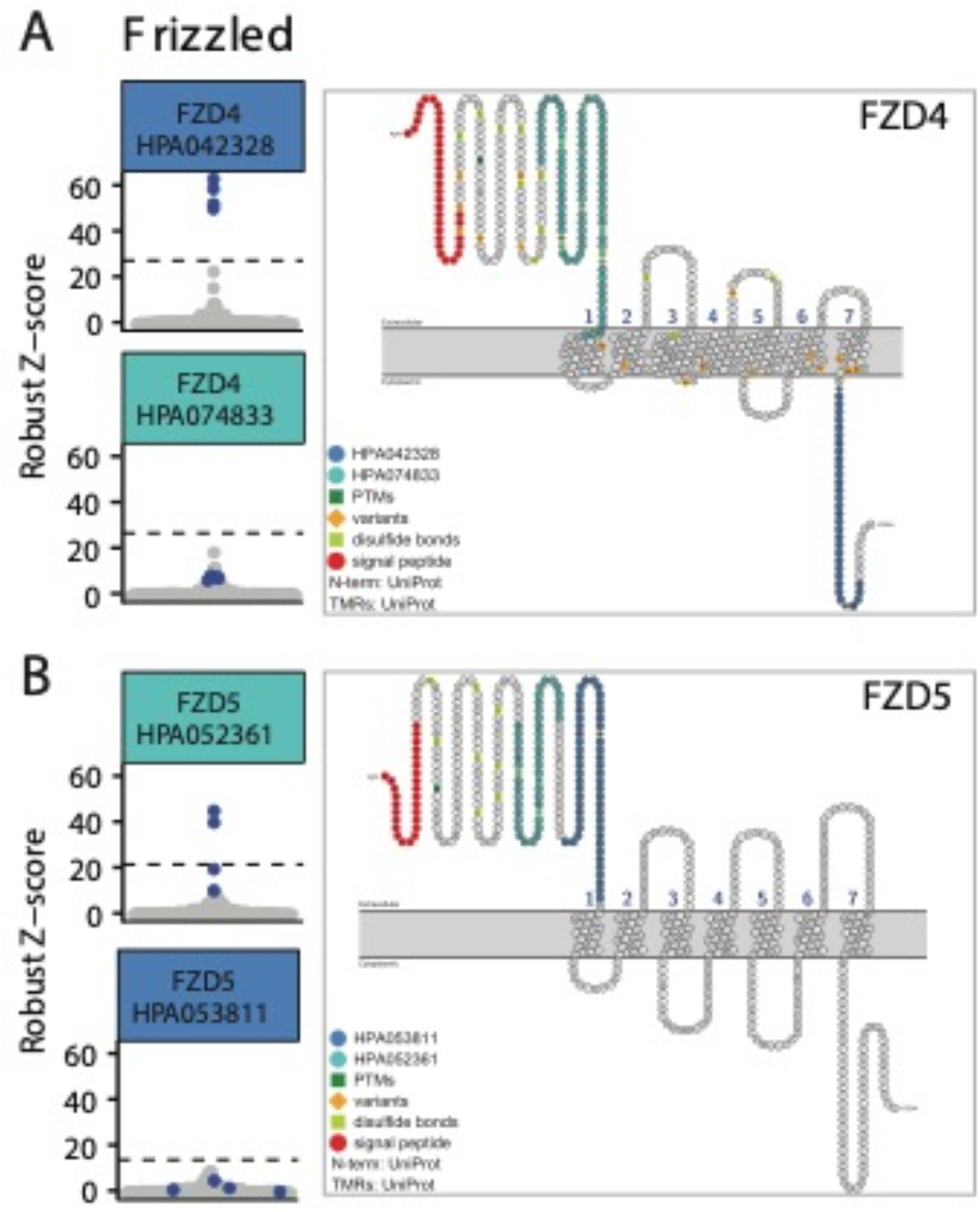
Detection of FZD GPCRs with paired antibodies. Left column: Beeswarm plots showing binding events for multiple Abs targeting the same FZD GPCR. Intended GPCR, blue dots; unintended GPCRs, grey dots. Dashed lines correspond to the selectivity cutoff for each HPA Ab. Color coding of HPA Ab ID corresponds to the color coding of the antigen in the snake plot diagram. Right column: Snake plot diagrams showing the antigen sequence used to generate the antibody on the entire protein sequence. Generated with Protter (*36*).

**Fig. S3.**
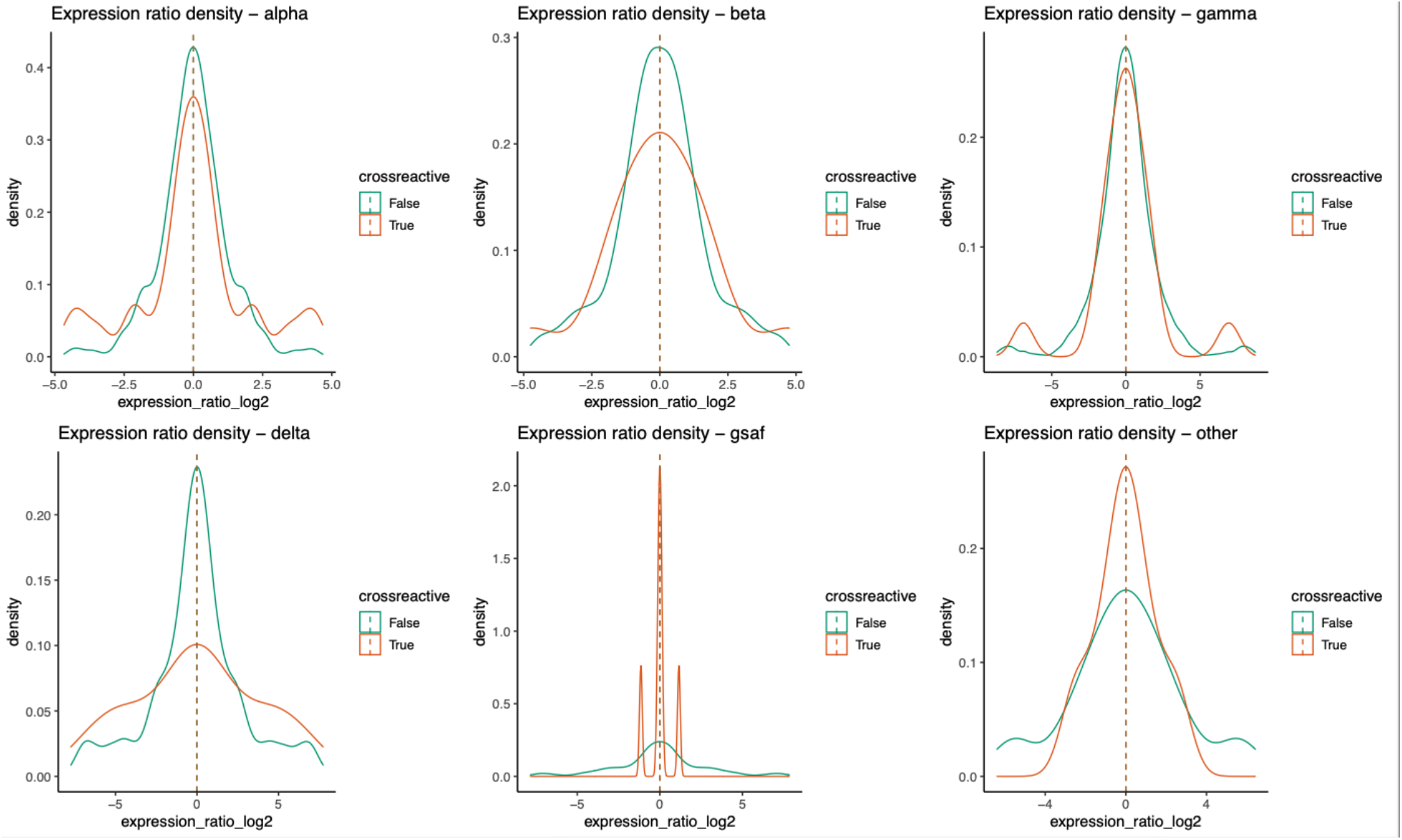
Expression ratio density plots per GPCR subfamily. The density plots illustrate the expression ratios between on- and off-target GPCRs for the different subfamilies. For the Abs in the rhodopsin alpha, gamma, and delta, and glutamate, adhesion, secretin, and frizzed (GSAF) subfamilies, there was a higher abundance of off-target GPCRs (orange) compared with on-target GPCRs (green).

**Fig. S4.**
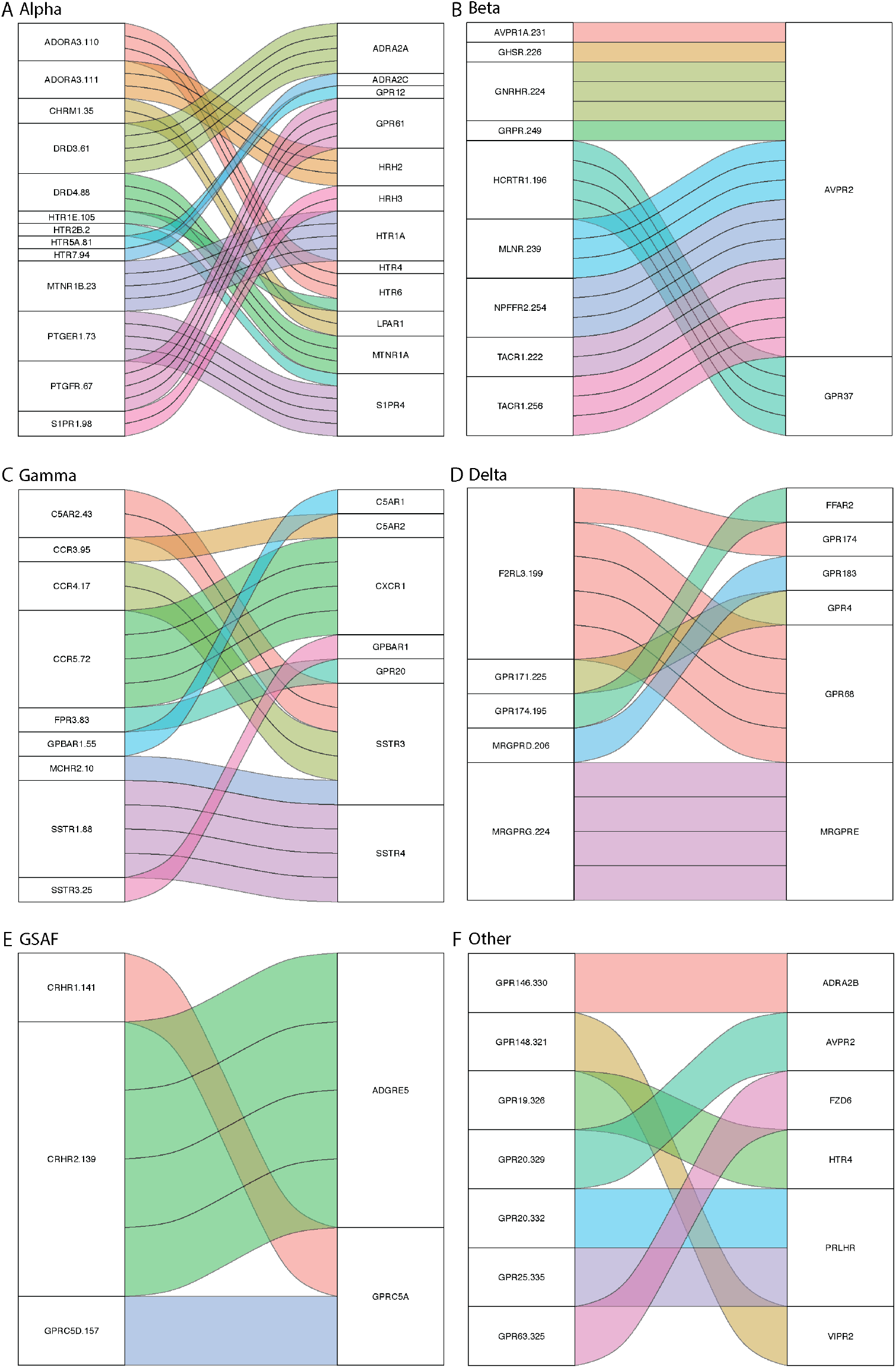
Summary of cross-reactivity per phylogenetic group. **(A-F)** Alluvial plots per GPCR subfamily (**A-D**: Rhodopsin alpha, beta, gamma, and delta, **E**: Glutamate, secretin, adhesion, frizzled (GSAF), **F**: Other) showing Abs on the left axis and their off-target GPCRs on the right axis. Axes are sorted alphabetically. Each line represents one GPCR-containing sample assayed with the Ab. The connections are colored by antibody.

**Fig. S5.**
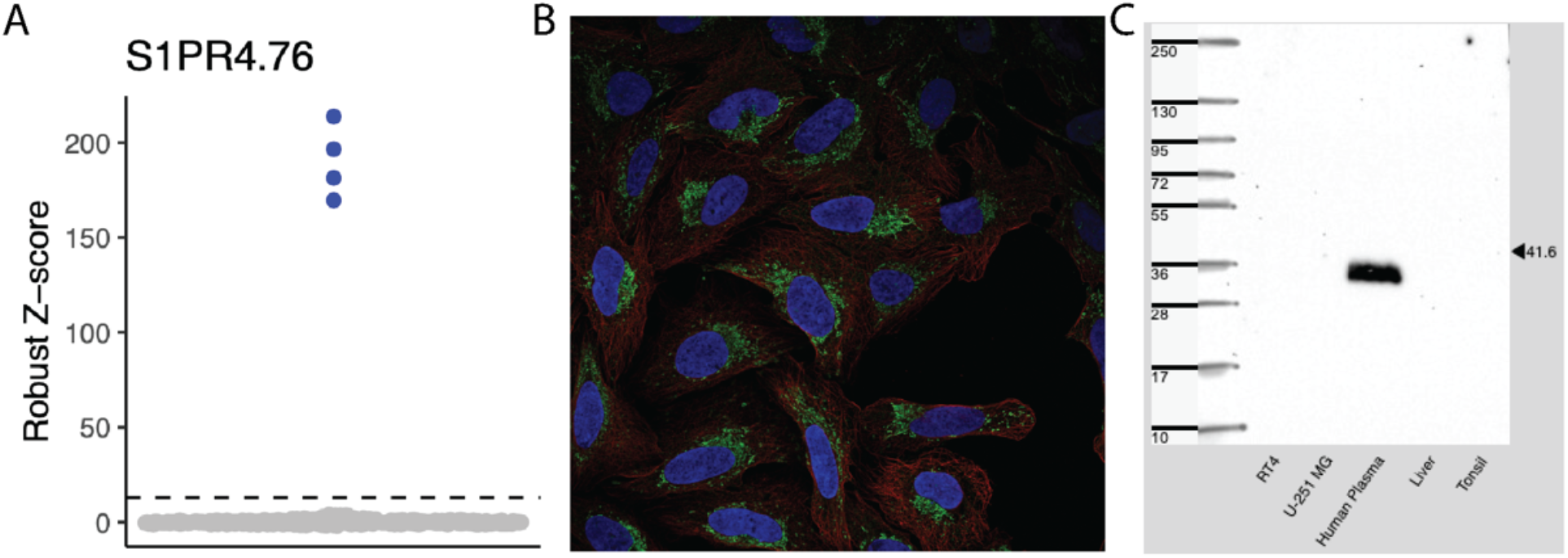
Utility of tested Abs in other assays. **(A)** An example of an on-target Ab used in other assays. The Ab HPA067232 selectively recognized the GPCR Sphingosine-1-phosphate receptor 4 (SP1R4) was validated by SBA. The results of the SBA assay are in line with the orthogonal validation performed by the HPA by immunofluorescence assay **(B)** and by Western blot **(C)**. Images from **(B)** and **(C)** are from the Human Protein Atlas [v21.proteinatlas.org].

## Supplementary tables

Tables are provided as separate files, and their legends can be found below.

Table S1. HPA antibodies and GPCRs. Listing all used HPA antibodies, their target GPCRs, and information about each GPCR and antigen. Performance of each antibody in the validation.

Table S2. GPCR expression test results per subfamily, per GPCR and for replicates of three select GPCRs.

Table S3. Structural predictions of Ab antigens.

